# A suspension technique for efficient large-scale cancer organoid culturing and perturbation screens

**DOI:** 10.1101/2021.10.23.464385

**Authors:** Stacey Price, Shriram Bhosle, Emanuel Gonçalves, Xiaodun Li, Dylan P McClurg, Syd Barthorpe, Alex Beck, Caitlin Hall, Howard Lightfoot, Luke Farrow, Rizwan Ansari, David A Jackson, Laura Allen, Kirsty Roberts, Charlotte Beaver, Hayley E Francies, Mathew J Garnett

## Abstract

Organoid cell culture methodologies are enabling the generation of cell models from healthy and diseased tissue. Patient-derived cancer organoids that recapitulate the genetic and histopathological diversity of patient tumours are being systematically generated, providing an opportunity to investigate novel cancer biology and therapeutic approaches. The use of organoid cultures for many applications, including genetic and chemical perturbation screens, is limited due to the technical demands and cost associated with their handling and propagation. Here we report and benchmark a suspension culture technique for cancer organoids which allows for the expansion of models to tens of millions of cells with increased efficiency in comparison to standard organoid culturing protocols. Using whole-genome DNA and RNA sequencing analyses, as well as medium-throughput drug sensitivity testing and genome-wide CRISPR-Cas9 screening, we demonstrate that cancer organoids grown as a suspension culture are genetically and phenotypically similar to their counterparts grown in standard conditions. This culture technique simplifies organoid cell culture and extends the range of organoid applications, including for routine use in large-scale perturbation screens.

## Introduction

Organoids are a three-dimensional long-term cell culture model grown in an extracellular matrix (ECM) with niche growth factors. They can be cryopreserved and remain genetically and phenotypically stable in culture^1,2^. Since the first reports of long-term patient-derived organoid models in 2011^3^, incremental technological advances have been made to improve cultures. These include intuitive next steps ranging from increasing the number of organs and diseases that can be modelled *in vitro*^1-14^, to the development of assays to interrogate a wide variety of biological questions^15-21^. Organoid cultures are accelerating both basic and translational research in a number of scientific disciplines from developmental biology to personalised cancer medicine^17,19,20,22^. In the setting of cancer, patient-derived tumour organoid cultures are valuable preclinical tools which complement existing models such as 2D cell lines and mouse models. They have been shown to recapitulate clinically relevant responses to therapy and are being evaluated as patient avatars to individualise patient treatment^17,23,24^.

Screens in 2D cancer cell lines have been broadly adopted to identify novel gene-drug interactions as well as drug targets to advance precision cancer medicine^25-30^. However, currently available cell line collections do not adequately represent the genomic complexity and clinical landscape of cancer^31-33^. The derivation of comprehensive panels of tumour organoid models are increasing this representation. High rates of success during derivation and the ability to derive from disease stages and treatment pathways that were once impossible or extremely difficult^1-3,7,11,34-36^, has allowed the community to increase and widen the number of cell models available, one such effort to develop models is the Human Cancer Models Initiative (https://www.sanger.ac.uk/collaboration/human-cancer-model-initiative-hcmi//, https://hcmi-searchable-catalog.nci.nih.gov/).

Organoid cultures can be used for many molecular and cellular biology experiments routinely applied to 2D culture. Nevertheless, there are practical limitations when working with organoid models. In comparison to 2D cell culture, the culturing of organoids is more expensive due to the requirement of an ECM and specialised growth medium. The rate of organoid growth, in general, is slower than traditional cancer cell lines. In addition, standard culture protocols embed the organoids within small polymerised domes of 80% - 100% ECM (protein concentration 6.4-12 mg/ml). While this technique is feasible for model derivation and when working with small cultures, when working with larger numbers of organoids this can be labour intensive and restrictive with respect to reagent costs and ergonomics. For example, to conduct a genome-wide CRISPR-Cas9 knock-out screen in technical triplicate would on average require up to fifteen 6-well plates. The challenges associated with efficiently growing organoids, particularly for large-scale phenotypic assays, currently limits their utility for some applications.

Here, we aimed to develop an alternative culture technique to improve the efficiency and handling of organoid cultures to facilitate their use in perturbation screens as well as other applications where large numbers of cells are required. We developed culture conditions using a low concentration of ECM (5%, protein concentration 0.4-0.6 mg/ml) that supports large-scale organoid expansion and long-term culturing. Similar approaches have been utilised previously for specific, short-term applications^9,12,14^, but importantly have not been assessed for long term culturing and expansion of models, nor has the impact on culture phenotypes been assessed. We show that in low percentage ECM conditions organoid cultures grow in suspension making handling easier and reducing costs through the decreased volumes of ECM required. Furthermore, we confirm that the genomic landscape and phenotypic screening results, including drug sensitivity testing and CRISPR-Cas9 genome-wide screening, are consistent between low ECM and standard conditions. This new culturing approach improves the efficiency of organoid culturing and facilitates their use in a wide range of applications including large-scale perturbation screens.

## Results

### A low ECM concentration organoid culturing technique

In order to address the practical, ergonomic and cost implications of culturing organoids at scale several alternative techniques were initially investigated and compared to standard culture protocols using 80% ECM. These included ECM (basement membrane extract-2 (BME-2)) droplets suspended in media and cultured in spinner flasks^22^, microcarrier beads in a 5% ECM solution and an ECM concentration gradient in ultra-low attachment plates. For all techniques tested, published organoid media recipes were used. Spinner flasks and microcarrier beads failed to support organoid growth and did not improve the ease of handling cultures. In contrast, lower percentages of ECM appeared to address the necessary requirements effectively (Table 1).

**Table 1.** Three methodologies tested for organoid culturing compared to standard organoid culturing protocols.

| Technique | Organoid Growth | Scalability | Time (benefit) | Cost (benefit) |
| --- | --- | --- | --- | --- |
| Standard Conditions 80% ECM | Yes | + | + | + |
| 5% ECM | Yes | +++ | +++ | +++ |
| Microcarriers | Yes | - | + | + |
| Spinner Flasks | No |  |  |  |

A gradient range of 0, 5, 20 and 50% ECM was trialled using the colorectal cancer organoid HCM-SANG-0266-C20 in standard and ultra-low attachment 6-well plates (details for organoid cultures are provided in Supplementary Information Table 1). Organoids were plated as single cells and over 5-7 days organoid formation was assessed. Ultra-low attachment plates supported the formation of typical organoid structures, whereas conventional cell culture plates led to the organoids adhering to the bottom of the plate and the loss of 3D structures (Supplementary Figure S1); all further optimisation experiments used ultra-low attachment plates. In 0% ECM some organoid formation was observed, but the culture was dominated by a loss of cell viability. For both 5 and 10% ECM, organoids grew as a suspension culture, attaching to small pieces of ECM that had polymerised in the organoid media. Concentrations beyond 20% ECM led to complete polymerisation of the ECM and organoid media forming a solid ECM layer, more akin to standard fully-polymerised dome culture conditions. The solid ECM layer formed using >20% ECM did not facilitate easier handling or significantly reduce the volume of ECM compared to standard conditions. Short-term organoid formation and growth was supported for one week in as little as 5% ECM in ultra-low attachment plates (Supplementary Figure S1). Based on these initial studies 5% ECM was taken forward for further evaluation.

We next evaluated whether 5% ECM supports the formation and expansion of a range of organoid models. A further 4 colorectal, 8 oesophageal and 5 pancreatic cancer organoid models were successfully cultured for over one week (n = 18 models), demonstrating the wider application of this approach in different tumour types (Supplementary Figure S1). Organoids when grown in 5% ECM generally appeared larger in comparison to standard 80% organoid culturing techniques, also visible by H&E staining (Figure 1a), possibly due to not being physically confined within a 20μl dome of ECM. The lack of confinement also led to an increased tendency for individual organoids to adhere together, again contributing to the formation of large organoids. Apart from being larger in size when grown in 5% ECM, no morphological differences were observed by H&E staining when compared to the counterpart models grown in standard conditions (Figure 1a). Furthermore, the proliferation marker Ki67, cytokeratin staining and p53 expression patterns by immunohistochemistry were consistent in both culture conditions.

**Figure 1.**
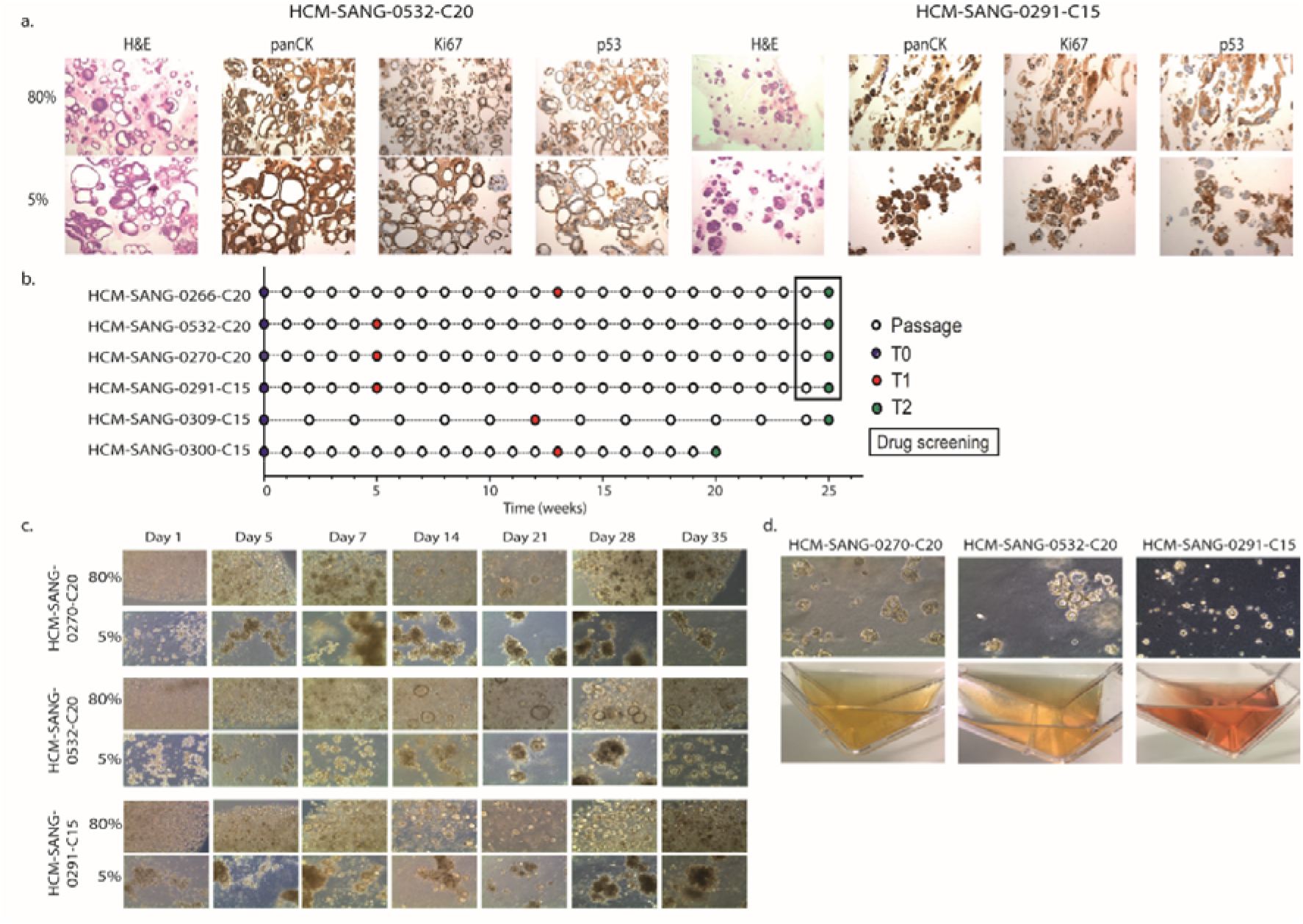
Alternative organoid culturing techniques and longitudinal study to assess 5% ECM culturing feasibility. a. Representative images of H&E staining and IHC of panCK (pan-cytokeratin), Ki67 and p53 in two organoids cultured in either 80% ECM domes or 5% ECM suspension. All images are 10X magnification. b. Study design comparing the 5% ECM with standard conditions. Samples were taken at the start (T0), after approximately 1 - 3 (T1) and 6 (T2) months. c. Representative images of three organoid models over the first 35 days of the study. d. Images of three organoid models growing in T75 flasks in 5% ECM. Organoid images are 10X magnification

In order for the 5% ECM technique to be applicable for large scale expansion of organoid models, prolonged culturing over multiple months and passages is desirable. To determine if the 5% ECM culture could support long-term organoid expansion, six models were continuously cultured longitudinally in parallel for up to six months in standard and 5% ECM conditions. During this time they were subjected to genomic and phenotypic characterisation to ensure the culture conditions had little to no impact on genomic evolution in culture, as well as observed drug and gene dependencies (Figure 1b). The appearance of these 5% cultures over the first 7 weeks further support the observation that organoids appear larger and are able to aggregate in low ECM conditions (Figure 1c). The 5% ECM suspension culture method provides technical benefits over standard organoid culturing protocols, including a decreased time requirement per passage, less physical handling of ECM to reduce injury risk due to repetitive motions, and lower volumes of ECM equating to a cost reduction. This technique can be scaled to use ultra-low adherent flasks (figure 1d), rather than multi-well plates, minimising incubator space requirements as well as reducing the risk of microbial and cross-model contamination.

### Genomic characterisation of organoids grown in 5% ECM suspension culture

*In vitro* disease models can evolve over time in culture^37^, and acquire new mutations through both cell intrinsic processes as well as extrinsic selective pressures such as culture conditions. In characterising the 5% ECM culture method it was important to ensure that these culture conditions were not directly contributing to or adversely influencing how the models evolve while in culture. In order to assess this, samples from the six models grown longitudinally (T0, T1, T2) in both 5% ECM and standard conditions (Figure 1b) underwent whole-genome sequencing (WGS) and RNA sequencing (RNAseq).

We began by comparing all single nucleotide polymorphisms (SNPs) identified in all samples using WGS. The variant allele fraction (VAF) of SNPs, irrespective of time point or culture condition, were highly correlated with organoids at T0 grown in standard conditions when considering all SNPs or only non-synonymous SNPs. (Figure 2a and Supplementary Figure S2). Furthermore, the total mutational burden in the six models remained consistent in both conditions over the six-month timeframe (Supplementary Figure S2), as well as the distribution and pattern of mutations across the genome (Supplementary Figure S3). A very high concordance of identified variants and insertions and deletions was observed in each culture condition at each time point; and in 4 out of the 6 models, concordance was greater than 90% at both time points. Additionally, this was corroborated by high jaccard index scores comparing variants for each time point; the Jaccard index is a measure of similarity between samples determined by comparing the intersection of two sample sets (Figure 2b). There were a small number of mutations in all models and at all timepoints that were exclusive to both 5% and 80% culture conditions. None of these mutations were in cancer driver genes and could represent sequencing artefacts, clonal selection of tumour sub-populations during culturing, or the acquisition of mutations due to different mutational processes operative in each of the culture conditions.

**Figure 2.**
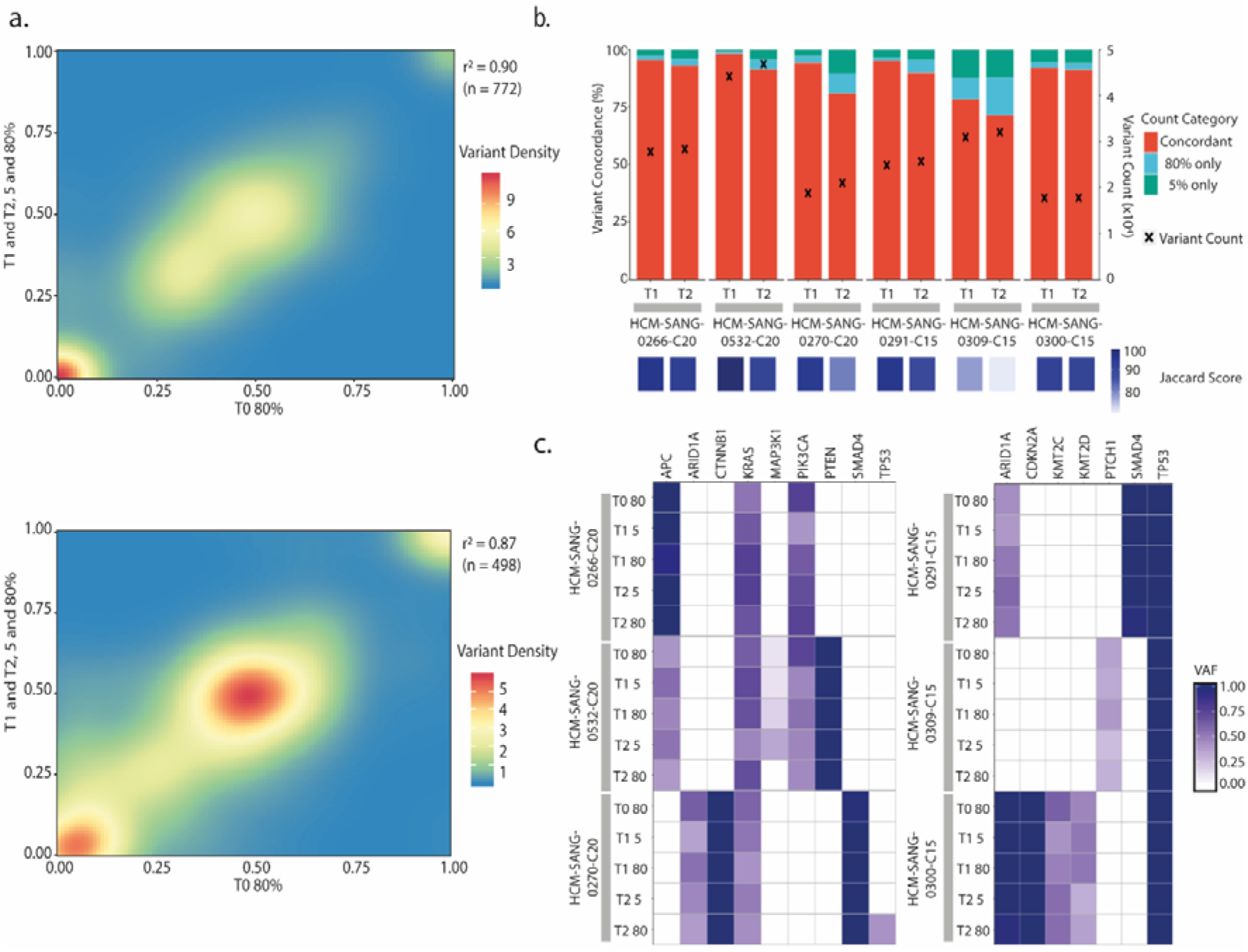
Genomic stability analysis. a. Correlation density plot of the VAF for all non-synonymous variants at T0 in standard conditions with all other time points in both culture conditions, for the three colon models (top) and three oesophageal models (bottom). b. Concordance of mutations with a VAF greater than 0.05 for each model with associated jaccard index scores, and total variant count considered at each time point. c. Intogen database filtered cancer-specific driver gene VAF heat maps for colon samples (left) and oesophageal samples (right).

We focused next on the presence of mutations in cancer driver genes known to have a pathogenic role in the respective cancer types. Notably, the identity and VAF of cancer driver mutations^38^ was retained between both standard conditions and 5% ECM after prolonged culturing (Figure 2c). For example, *KRAS* mutations were observed in all colon models under both conditions and time-points, and as expected the oesophageal adenocarcinoma models all have *TP53* mutations irrespective of growth conditions. There was one example where a missense driver mutation in *TP53* (17:7577548:C:T, VAF 0.41), was acquired in HCM-SANG-0270-C20 at T2 under standard conditions, most likely reflecting selection of a rare sub-clone present at T0 but below detection. We also observed loss of *MAP3K1* in HCM-SANG-0532-C20 at T2 in 80% ECM, but this is retained in the T2 5% ECM sample (Figure 2c). Taken together these data indicate that the cultures do not have any global mutational alterations or divergence in the representation of driver mutations when cultured up to 6 months in 5% ECM.

Many cancers are driven by copy number alterations that contribute to tumour phenotypes. Using the mean logR copy number of all genes, we observed a high correlation within samples from each model (Supplementary Figure 4). Furthermore, the global pattern of copy number alterations at T0 and T2 was shown to be consistent across time points in both culture conditions. Importantly, the ploidy remained consistent across all conditions from a particular model, and when focussing on tissue-specific copy number driven cancer genes the vast majority have a consistent copy number for all time points and culture conditions (Figure 3a)^39,40^. Furthermore, clustering of all samples at T0, T1 and T2 by RNA-sequencing derived gene expression showed all samples clustering by model (Supplementary Figure S4). Unsupervised hierarchical clustering of the 3,000 most differentially expressed genes demonstrated that, in the majority of models, samples clustered first by organoid model, and then by time point (Figure 3b). This demonstrates that the time point at which the RNA was harvested had more impact on gene expression profiles than the culture condition.

**Figure 3.**
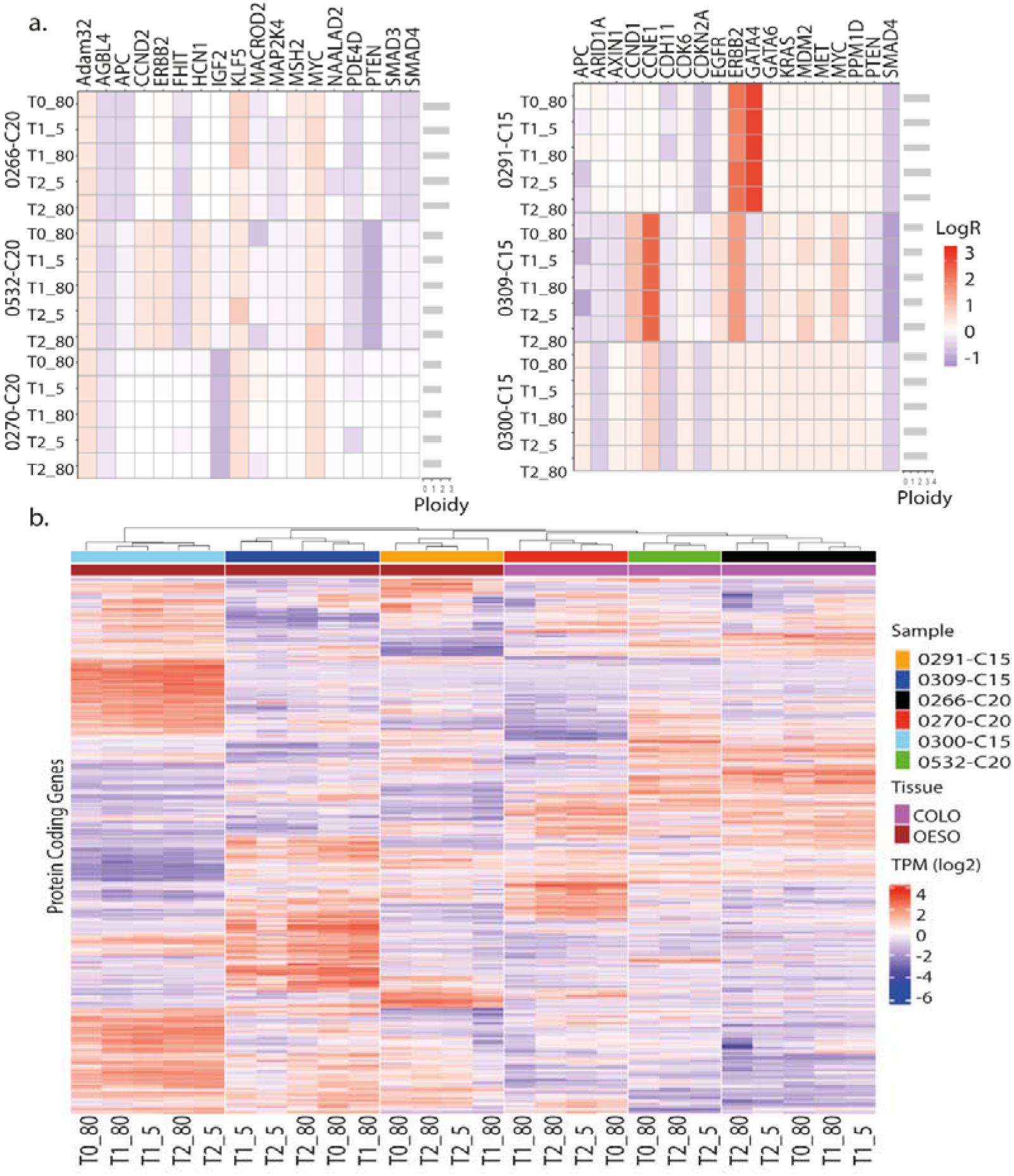
Copy number variation and gene expression in 5% ECM and standard 80% ECM culture conditions. a. Tile plot showing the LogR values of copy number altered tissue-specific cancer driver genes. b. Unsupervised hierarchical clustering of log TPM values, for the 3,000 most differentially expressed genes. The colour scale shows square root of standardised log_2_ TPM values scaled by the standard deviation and centred to the mean.

Collectively, our WGS and RNAseq analysis confirms that models cultured in 5% ECM are genetically and transcriptionally virtually indistinguishable to those cultured in standard 80% ECM droplets.

### High-throughput drug sensitivity testing of organoids grown in low ECM culture

Drug sensitivity testing in cancer cells is an important application in drug development and so we sought to determine whether long term culture in 5% ECM conditions led to changes in sensitivity. Using established protocols for organoid drug testing^2,6,10,13^, the sensitivity of four organoid models to 72 anti-cancer drugs was compared after propagation for up to six-months (T2) in standard or 5% ECM cultures conditions. Drug sensitivity was measured as 1 – area under the dose response curve (1-AUC) determined from a 7-point half-log dilution series encompassing a 1,000-fold concentration range. The compounds assayed included FDA-approved oncology drugs, compounds in clinical development, investigational compounds to a wide range of oncology targets, and also included chemotherapeutic agents. As expected, we observed a range of sensitivities to different drugs tested in the organoids (Figure 4a and Supplementary Information Table 2). Notably, the sensitivity to the drugs of the four organoids grown for 6 months in 5% or 80% ECM conditions were highly correlated (Pearson correlation > 0.94).

**Figure 4.**
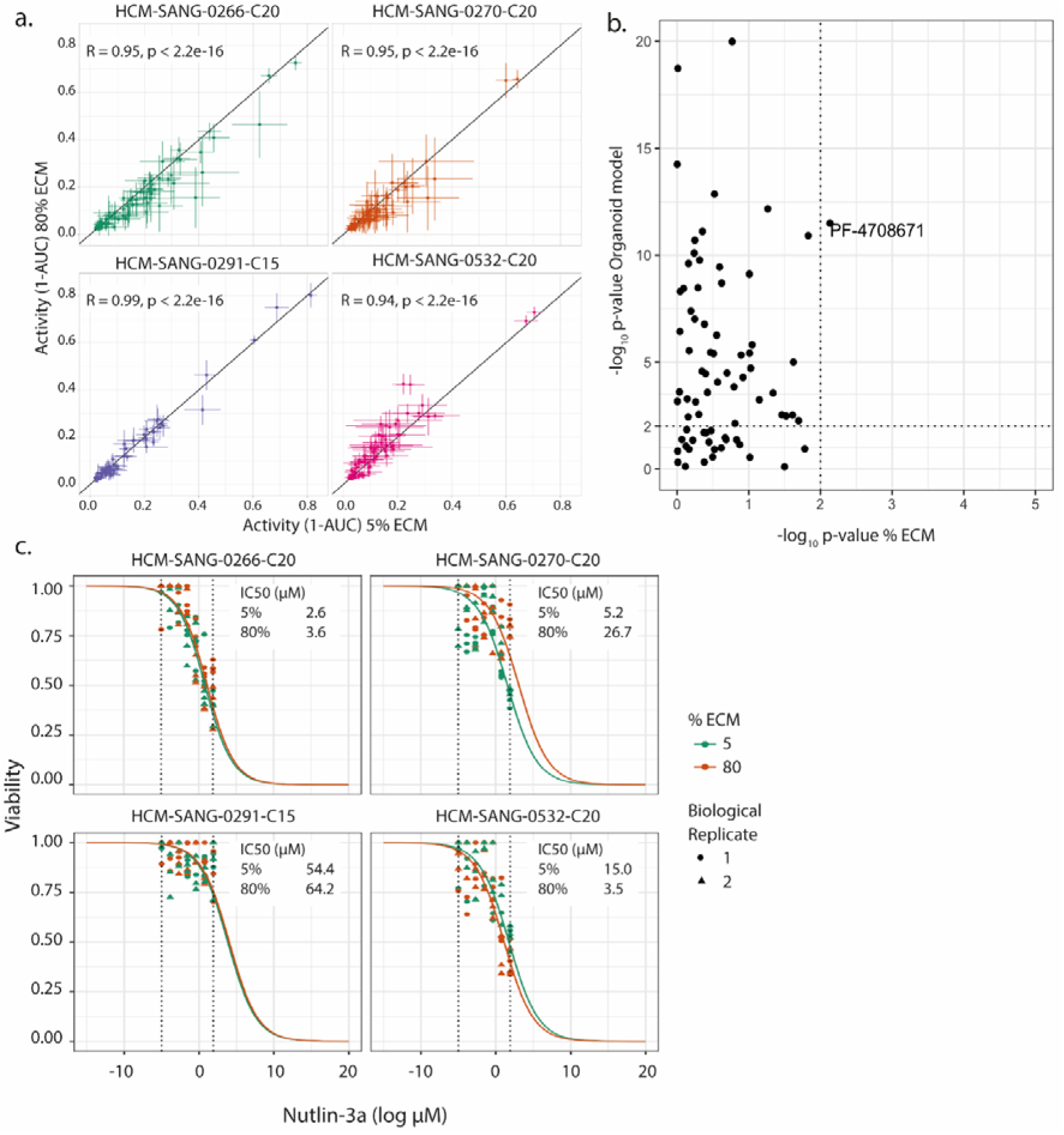
Drug response in standard vs 5% ECM conditions. a. Sensitivity of four organoids to 72 anti-cancer drugs in 5% ECM (x-axis) and 80% ECM (y-axis). Each data point has error bars which represent the 95% confidence intervals using the t-statistic (n= 5-12 from two biological replicates). b. ANOVA output of drug response and dependence on either culture condition or organoid model. Dashed lines are thresholds for statistical significance (p < 0.01). c. Representative dose response curves for the four organoid models when treated with Nutlin-3a. Cells cultured in 5% ECM prior to drug screening are coloured orange and 80% ECM are coloured blue. Biological replicates are shown by different shapes and dotted lines indicate the Nutlin-3a concentration range.

An analysis of variance (ANOVA) was performed to systematically identify whether the model or culture condition had the greatest impact on sensitivity of the organoids to each of the 72 individual drugs. The organoid model had a significant effect on the sensitivity to many drugs (n =54 drugs at p-value threshold 0.01), whereas for nearly all drugs the culture condition had no significant impact (Figure 4b). The only exception was the S6K1 inhibitor PF4708671, where 5% ECM was significantly associated with reduced sensitivity in HCM-SANG-0532-C20. However, this significance is marginal and was only observed in a single model suggesting this might be due to an outlier data point (Supplementary Figure S5).

Known biomarkers of drug sensitivity were observed in organoid models cultured in both conditions. For example, a *TP53* wild-type colon cancer organoid HCM-SANG-0266-C20 was sensitive to the MDM2 inhibitor Nutlin-3a, whereas a *TP53* mutant oesophageal organoid HCM-SANG-0291-C15 exhibited little to no response to treatment; a model was deemed sensitive if the IC_50_ was within the screening concentration tested (Figure 4c). The two additional *TP53* WT models did not show sensitivity. All *KRAS* mutant colon cancer models were insensitive to EGFR inhibition (Supplementary Figures S6-8). We observed differential sensitivity to the ERK inhibitor SCH77298 in HCM-SANG-0266-C20 due to culture conditions, which appeared to be the result of a single outlier replicate data point in the 80% ECM condition (Supplementary Figures S6 and S10).

Taken together, these data indicate that long-term culturing in 5% ECM conditions prior to screening does not impact on the response of organoids models to a wide range of different anti-cancer drugs.

### CRISPR-Cas9 whole-genome knockout in 5% ECM culture

Another important application in oncology is the use of genetic perturbation approaches, such as loss-of-function CRISPR-Cas9 screens, to identify dependencies across tumour types. To further investigate whether phenotypic response is stable in 5% ECM conditions, a whole-genome CRISPR-Cas9 knockout screen targeting 18,009 genes (Yusa v1.1 library)^28^ was performed in the colorectal cancer organoid model HCM-SANG-0273-C18. To minimise variation, prior to transduction with the sgRNA library the organoids were cultured under standard 80% conditions, and following sgRNA transduction they were cultured for 3 weeks to allow drop-out of sgRNAs targeting fitness genes in either 5% or 80% ECM conditions (arm 1 and arm 2 respectively, Figure 5a and Supplementary Information Table 3).

**Figure 5.**
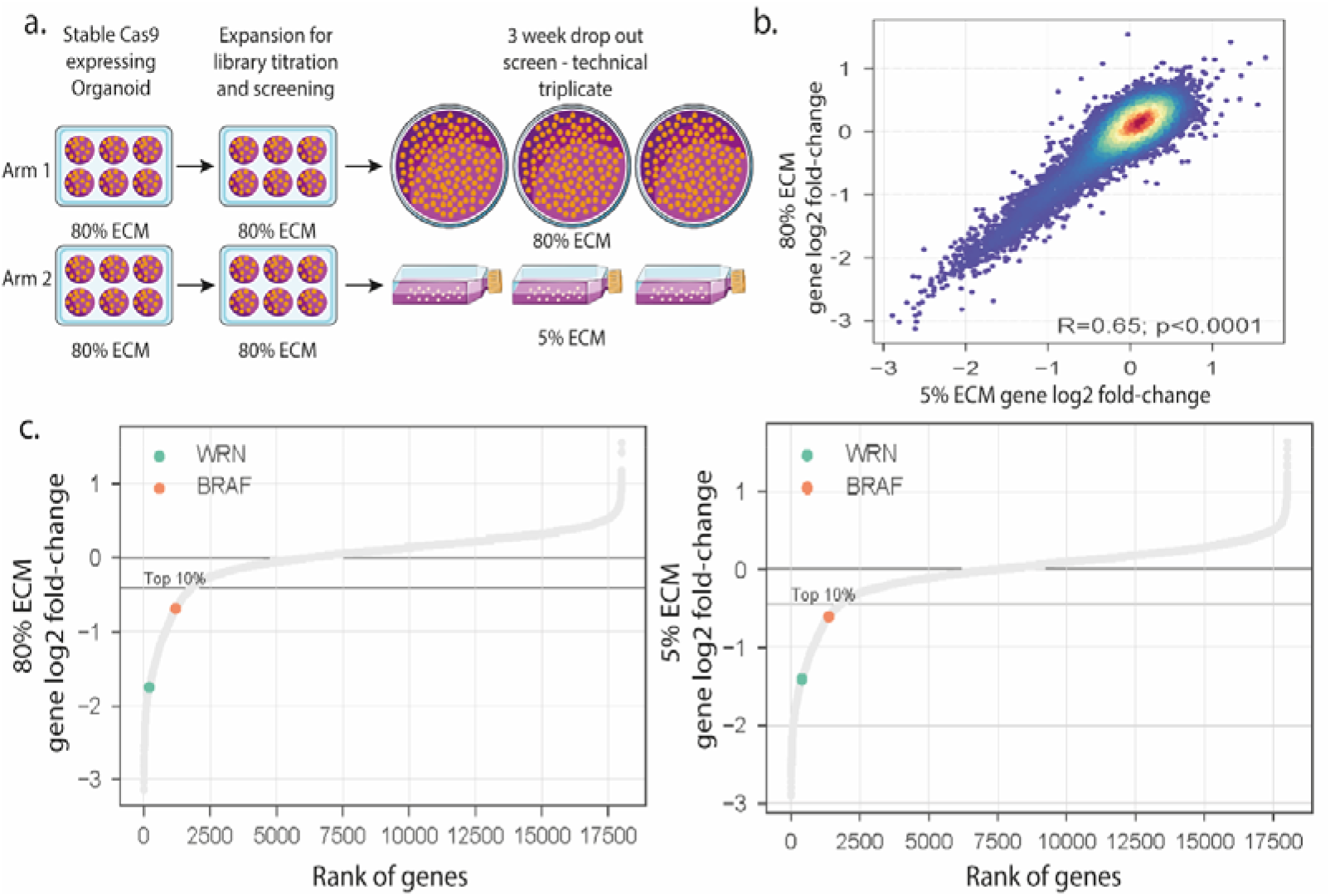
Comparison CRISPR screening in standard versus low ECM conditions. a. Schematic of the two experimental arms of the whole-genome CRISPR-Cas9 screen in HCM-SANG-0273-C18. b. Spearman’s Rho correlation between 5% and 80% ECM CRISPR-Cas9 screens at the gene level. c. Genes ranked by their log fold change from both experimental arms. WRN and BRAF are highlighted in the top 10% of dependencies.

As a measure of screen quality, the recall of known essential genes (area under the recall curve (AURC) = 0.93-0.94) was similar for both conditions, and the AURC values are equivalent to what is typically observed for screens performed in 2D cancer cell lines^28,30^ (Supplementary Figure S11). The distribution of the sgRNA and gene level fold changes were centred at 0 (Supplementary Figure S11) indicating that coverage of the library at the sgRNA and gene level was maintained in both culture conditions, confirming no unexpected loss of transduced cells during the three-week assay. There was a strong positive correlation between gene log-fold change values at both conditions (Pearson correlation R = 0.65), confirming that the identified fitness genes were consistent between culture conditions, particularly among the strong dependencies (Figure 5b and Supplementary Figure S11).

We were able to confirm known gene dependencies in HCM-SANG-0273-C18. For example, this model is a microsatellite unstable (MSI) colorectal cancer model harbouring a *BRAF* V600E mutation and, as expected, knockout of *BRAF* led to a strong loss of viability in both 80% and 5% conditions (top 10% dependencies, Figure 5c). Furthermore, knockout of *WRN* led to loss of viability in both conditions in keeping with its recent synthetic-lethal association with microsatellite instability^28^. This data indicates that CRISPR-Cas9 knockout screens can be performed in 5% ECM conditions without loss of signal while reducing the cost and, importantly, making these types of screens practical, especially when large numbers of models are screened.

## Discussion

We have developed a culture methodology that increases the efficiency of growing organoid cultures at scale while retaining genomic and phenotypic stability relative to standard organoid culture conditions. We have extensively benchmarked our approach using multiple cancer organoid cultures from three different cancer types, monitoring global and disease relevant molecular changes and phenotypes including drug sensitivity and genetic dependencies. Moreover, we provide direct evidence that it is possible to readily propagate organoid cultures in sufficient numbers making them suitable for systematic perturbations assays. Collectively, these results demonstrate the utility of using 5% ECM organoid cultures for many applications.

Current approaches for propagating organoids are time consuming, labour intensive and ergonomically challenging, and in practice limits the application of organoids for high-throughput and screening approaches. Using 5% ECM, we can successfully stably grow organoids in a suspension-like culture for prolonged periods. Notably, using low ECM reduces the cost of ECM by approximately 30-40% relative to the 80% ECM droplet equivalent and significantly improves manual handling. Previous studies have used ECM pre-coated plates or low ECM cultures, for short-term applications up to 5 days, including small scale drug testing^9,12,14^. However, unlike the study shown here, they had not evaluated the suitability of low ECM for long-term cultures, or for large-scale drug testing or CRISPR-Cas9 screens. Furthermore, the impact on important tumour cell characteristics including morphology and histology, mutations, copy number alterations, drug sensitivity, and genetic dependencies has not been evaluated. Thus, our study is the first to develop and benchmark this method suitable for key biological applications in cancer biology and high-throughput screening.

This new culture method provides a robust alternative technique for cost effective and easier expansion of organoids for high-throughput perturbation screens. This technique has not been used for the derivation of organoids from patient tissue; from our experience, the main benefit of this technique comes from when working with large numbers of cells, which is not the case at the point of model derivation. In conclusion, the methodology described here enables the efficient use of three-dimensional organoid models for perturbation screens at reduced cost, and has the potential to expand the range of applications accessible using organoid culture models and increase their application in cancer research.

## Methods

### Organoid Culture

All organoids were derived as part of the Human Cancer Model Initiative and model details are available from the Cell Model Passport (https://cellmodelpassports.sanger.ac.uk/).

For standard droplet culture, organoids were suspended in 20μl 80:20 ECM:Media droplets with a protein concentration of between 6.4-9.6mg/ml (BME-2, Amsbio 3533-005-02) on pre-warmed 6-well plates using previously published media recipes^2,10^. The droplets were incubated at 37 °C for 20 minutes before the addition of 2ml media to each well. For the 5% ECM technique, organoids are mixed with the same number of cells and volume of media as would have been used in the droplet condition, but with the addition of 5% ECM (e.g. for 1 well of a 6-well plate take 2ml media and 100μl BME-2) this suspension is mixed and immediately transferred to an ultra-low adherent plate or flask and placed into an incubator.

### Histology and IHC

Following BME-2 dissociation, organoids were gently pelleted and fixed in 10% neutral formalin before paraffin embedding and sectioning. Paraffin embedded sections of 3.5⍰μm were stained by a Bond Max autostainer according to the manufacturer’s instruction (Leica Microsystems). Primary antibodies cytokeratin (AE1/AE3, 1:100, Dako), Ki67 (8D5, 1:400, Cell Signalling Technology), and p53 (D07, 1:50, Leica) were applied with negative controls as previously described^2^.

### Whole genome DNA sequencing and analysis

Whole genome 150 base paired-end sequencing reads were generated using Illumina HiSeq X Ten platform. Reads were aligned using Burrows-Wheeler Alignment (BWA-MEM) tool^41^. PCR duplicates, unmapped and non-uniquely mapped reads were filtered out before downstream analysis. Single base substitutions and indels were identified using CaVEMan^42^ and cgpPindel^43^ respectively. Germline variants and technology-specific artefacts were removed by filtering against a matched normal blood sample and the panel of 100 unrelated normal samples (ftp://ftp.sanger.ac.uk/pub/cancer/dockstore/human/SNV_INDEL_ref_GRCh37d5.tar.gz). Additional post-processing filters were applied using in house post-processing tool cgpCaVEManPostProcessing (https://github.com/cancerit), variants sites that were flagged as ‘PASS’ were considered for further downstream analysis. Unbiased analysis of mutant and wild-type reads found at the loci of the base substitutions and indels were assessed across the related samples using vafCorrect^44^.

Copy number variant (CNV) analysis was performed using ASCAT^45^ algorithm wrapper ascatNgs^46^ specifically designed to process next-generation sequencing data. Copy number segments overlapping with summary intervals were merged and mean logR of these segments was assigned to a given summary interval and to the underlying genes in that interval, copy number states of these genes were used for downstream analysis. Driver genes with diploid status (logR = 0) were removed from the copy number tile plot.

### RNA sequencing and analysis

Paired-end transcriptome reads were quality filtered and mapped to GRCh37 (ensemble build 75) using STAR-v2.5.0c^47^ with a standard set of parameters (https://github.com/cancerit/cgpRna). Resulting bam files were processed to get per gene read count data using HTSeq 0.7.2. We calculated TPM (Transcripts per Million) values using the count and transcript length data for further downstream analysis. Only ‘protein coding’ genes (22,810) were considered for downstream QC and filtering steps. We used ‘filterByExpr’ (EgdeR)^48^ function with cutoff of 10 and 100 for *min*.*count* and *min*.*total*.*count* respectively with *min*.*prop* cutoff of 90%, resulting filtered genes (15,793) were used for sample level correlation analysis. Gene level clustering was performed on the top 8,500 genes ranked by variance.

### Drug sensitivity screening

Organoid drug sensitivity testing against 72 compounds was conducted in technical triplicate and biological duplicate. Formed organoids were seeded into 384 well plates onto a layer of ECM, and treated with 72 compounds over 3 days^2^. Dose response curves were fitted to experimental data using a non-linear mixed effects model^49^. The organoid model, the % BME media condition, and the drug treatment were included as random effects. Fitted curves with an RMSE of greater than 0.3 were removed from the data. Compound activity was calculated as 1 - AUC, where the AUC is the area under the curve within the screened dose range, thus the activity range is measured from 0 (no activity) to a maximum of 1. Analysis of variance was used to assess the effect of the organoid model and ECM condition on the activity of each compound, i.e. a linear model with the form: *compound ∼ % ECM + organoid model*.

### Whole-genome CRISPR knockout screening

Stable Cas9-expressing lines were generated using an overnight lentiviral transduction conducted in organoid media with 2.5μM Y-27632 and no ECM, plated the following day and antibiotic selection at day six. sgRNA library transduction was performed at 100X coverage of the Human CRISPR Library Yusa v.1.1^28^ with an MOI of 0.3, and following library transduction lines were cultured for 3 weeks. All plasmids and sample processing post library harvest were as described^28^.

### CRISPR Screening Analysis

CRISPR-Cas9 screen analysis was performed similarly to Goncalves et al, 2020^50^. Briefly, these started from sgRNA read count matrices. For each sample the number of sgRNAs with at least 10 counts was calculated to evaluate the library representation. Read counts were normalised to reads per million within each sample. Log2 fold-changes were calculated compared to the plasmid DNA (pDNA). Lastly, gene-level fold-changes were calculated by taking the mean fold-change of all targeting sgRNAs. Replicates were merged by averaging the gene-level fold-changes. Recall curves of essential and non-essential genes^30,51^ are estimated by ranking all the genes ascendingly according to their gene-level fold-change and the cumulative distribution is calculated. This is then summarized by estimating the area under the recall curve, where areas over 0.5 (random expectation) represent enrichments towards negative fold-changes, and areas lower than 0.5 represent enrichment towards positive fold-changes.

## Data availability

The whole genome DNA sequencing and RNA sequencing datasets are available in the EGA repository [accession numbers: EGAS00001003538, EGAD00001007971]. Details for organoid cultures are available through the Cell Model Passports database (https://cellmodelpassports.sanger.ac.uk/). All other data generated or analysed during this study are included in this published article and its Supplementary Information files.

## Acknowledgements

We thank Rebecca Fitzgerald for her assistance accessing samples and performing IHC. The laboratory of R.C.F. is funded by a Core Programme Grant from the Medical Research Council (RG84369). This research was funded in whole, or in part, by the Wellcome Trust Grant 206194 and 108413/A/15/D. For the purpose of Open Access, the author has applied a CC BY public copyright licence to any Author Accepted Manuscript version arising from this submission.

## Author contributions

S.P., H.F. and M.G. contributed to project conceptualization and design of the work. S.P led data acquisition and analysis with contributions from S.B., E.G., X.L., D.McC., Sy.B., A.B., C.H., H.L., L.F., R.A., D.A.J., K.R. and L.A. M.G., H.F., S.P., Sy.B., A.B., and C.B. contributed to project management. S.P., H.F. and M.G. drafted the manuscript. All authors approved the submitted manuscript.

## Competing interests

M.J.G. has received research funding from GlaxoSmithKline, Astex Pharmaceuticals and AstraZeneca, and is co-Founder and Chief Scientific Officer for Mosaic Therapeutics. M.J.G., S.P. and H.F. hold a patent application for the low ECM organoid culturing methods described in this study (GB2009808.3). All other authors declare no competing interests.

