## Supplementary Information for "A suspension technique for efficient large-scale cancer organoid culturing and perturbation screens"

**Supplementary Data**

**Supplementary Table S1:** Details of organoid models used in this study including their tissue type, cancer type, microsatellite instability status (MSI), American Type Culture Collection product number, and links to the respective Cell Model Passports Database cell model page.

**Supplementary Table S2**: Results from drug sensitivity testing of organoid cultures grown in 5% ECM and standard 80% ECM conditions.

**Supplementary Table S3**: Results from genome-wide CRISPR-Cas9 screens in organoids grown in 5% ECM and standard 80% ECM conditions. Gene-level log-fold change values are provided for each of three replicates in each condition.

**Supplementary Figures**

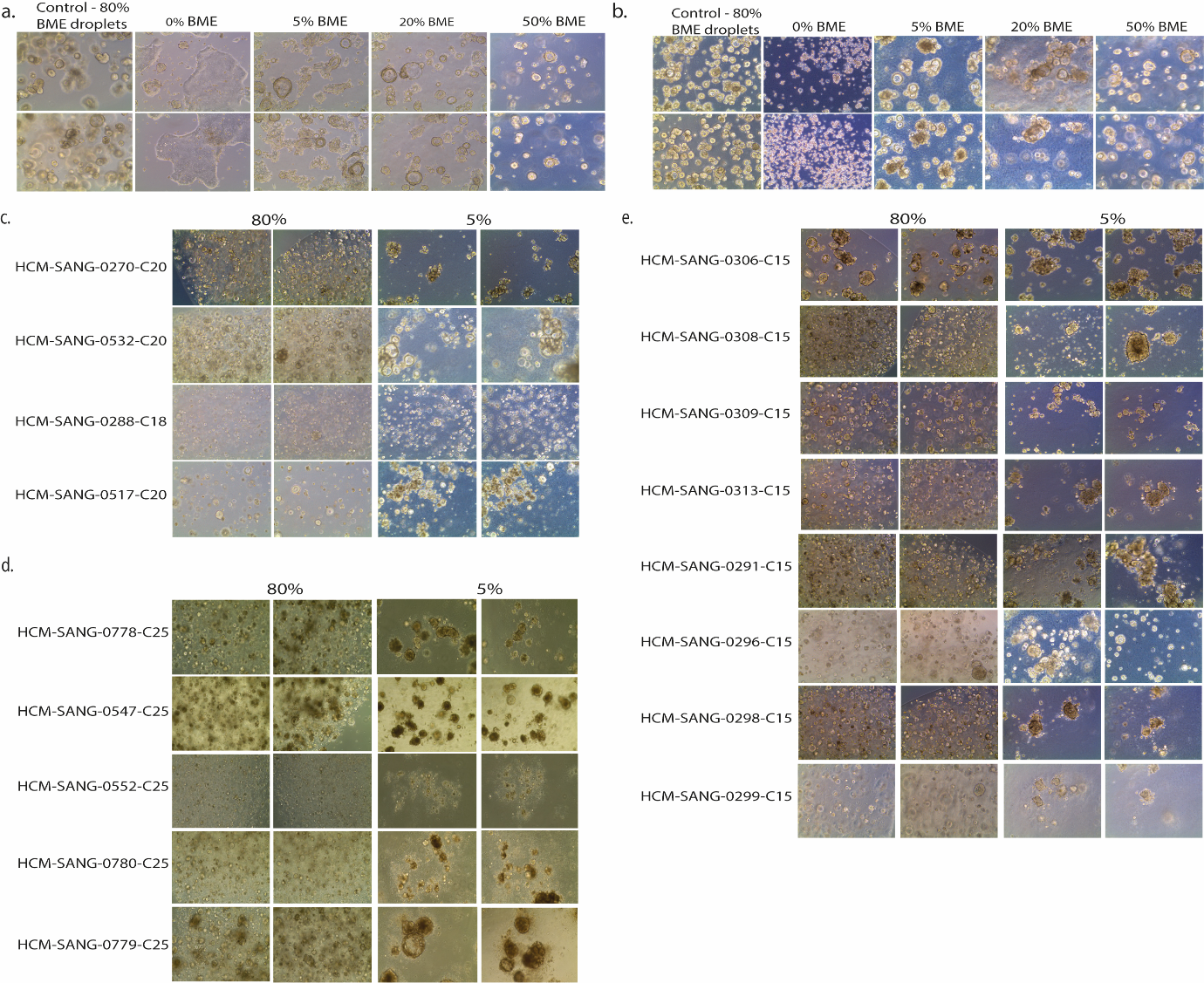

**Figure S1.** *a. and b. Representative images of HCM-SANG-0266-C20 7 days post seeding as single cells using a titration of ECM in a. standard tissue culture or b. ultra-low attachment 6-well plates. c., d. and e. Representative images of c. 4 colorectal cancer organoids, d. 5 pancreatic cancer organoids, and e. 8 oesophageal cancer organoids 3-7 days post seeding as single cells in either 80% ECM droplets (left) or 5% ECM (right) in ultra-low adherence plates.*

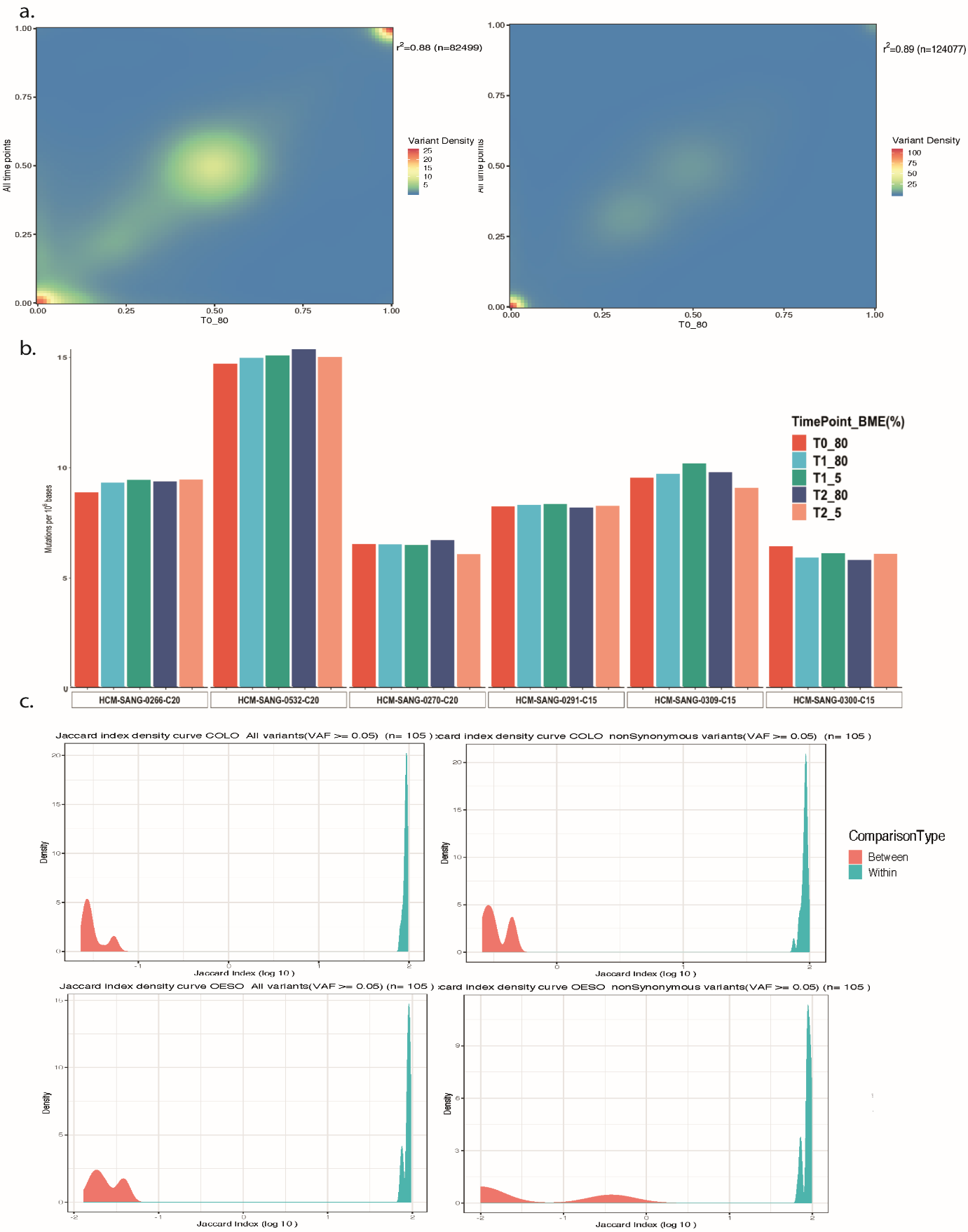

**Figure S2.** *a. Correlation density plot for 3 colon samples (left) and 3 oesophageal samples (right) comparing the variant allele frequency (VAF) for all variants at T0 with all other time points. b. Average number of mutations per million bases, grouped by model and coloured by time point and culture condition. c. Density plots for the log_10_ jaccard index when comparing within and between different models; all variants on the left and non-synonymous on the right.*

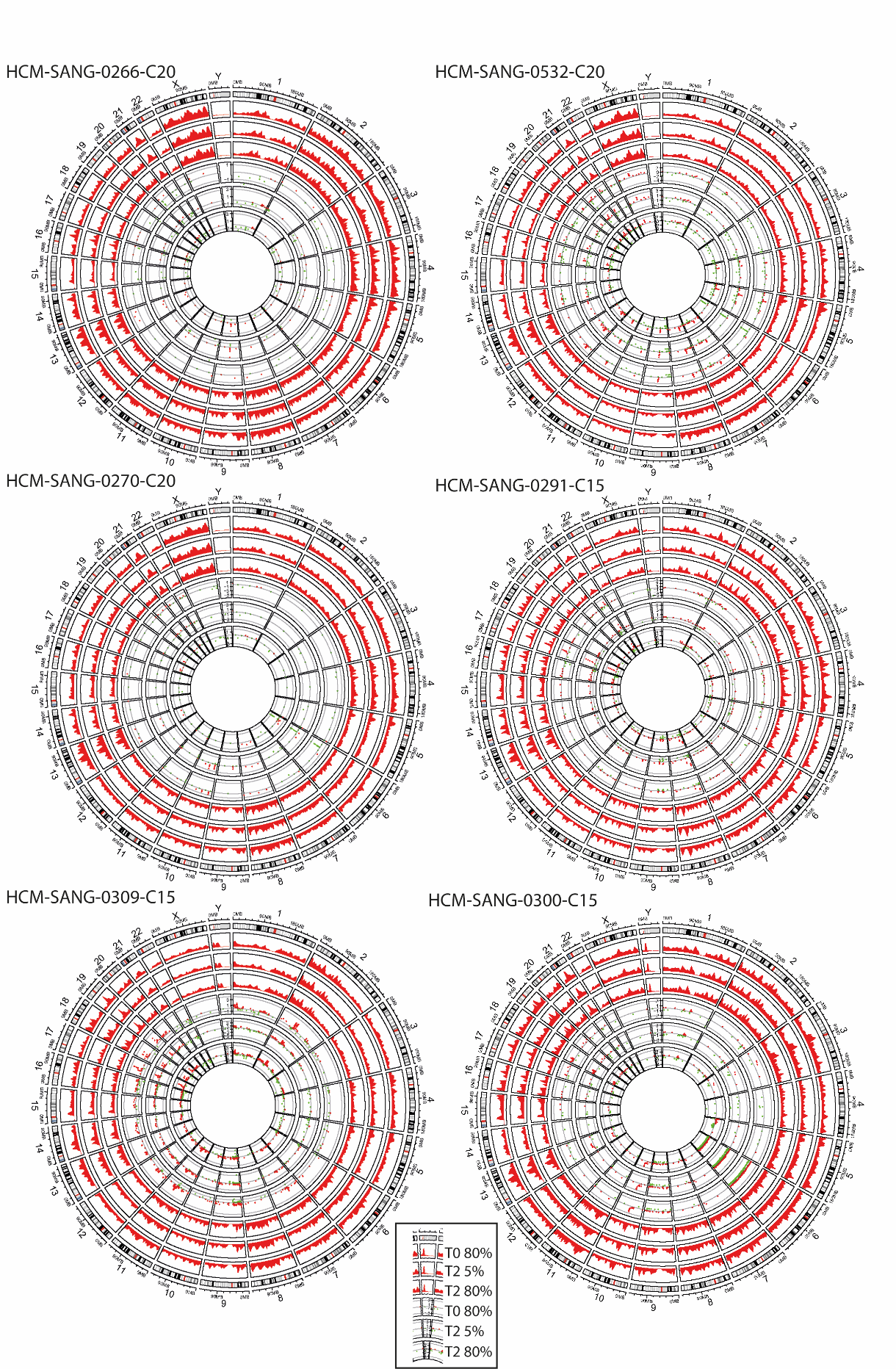

**Figure S3.** *Circos plots for 6 organoids. The outer 3 tracks show the distribution of variants across the genome in a 5mb window at T0 and T2 (5% and 80% BME), and in the inner 3 tracks the logR copy number at T0 and T2 (5% and 80% BME).*

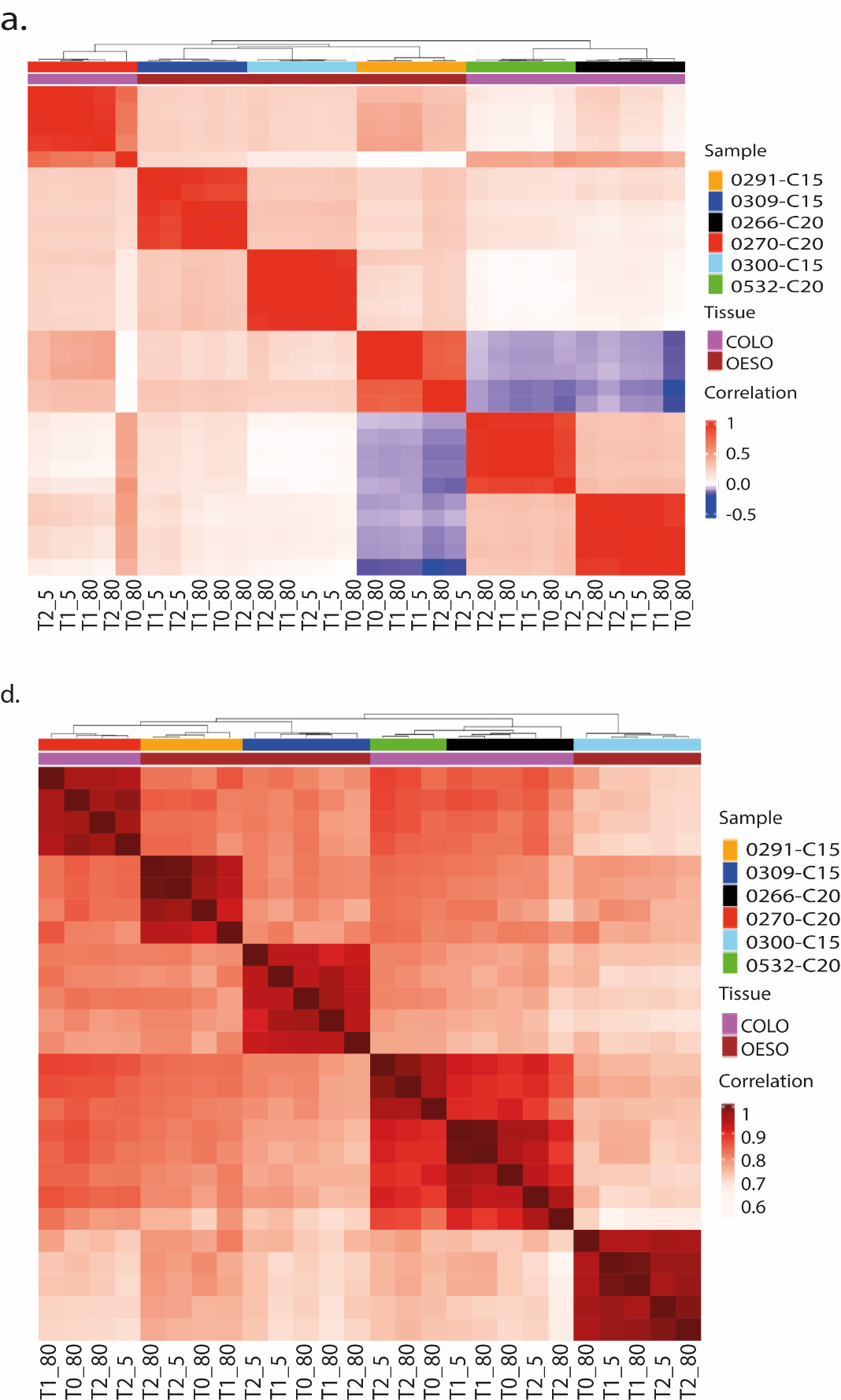

**Figure S4.** *a. Heatmap for all samples showing the logR copy number of all genes in six organoids at different timepoints (T0, T1 and T2) and culture conditions (5% or 80%). b. Heatmap showing the correlation of the gene expression of 12,228 protein-coding genes.*

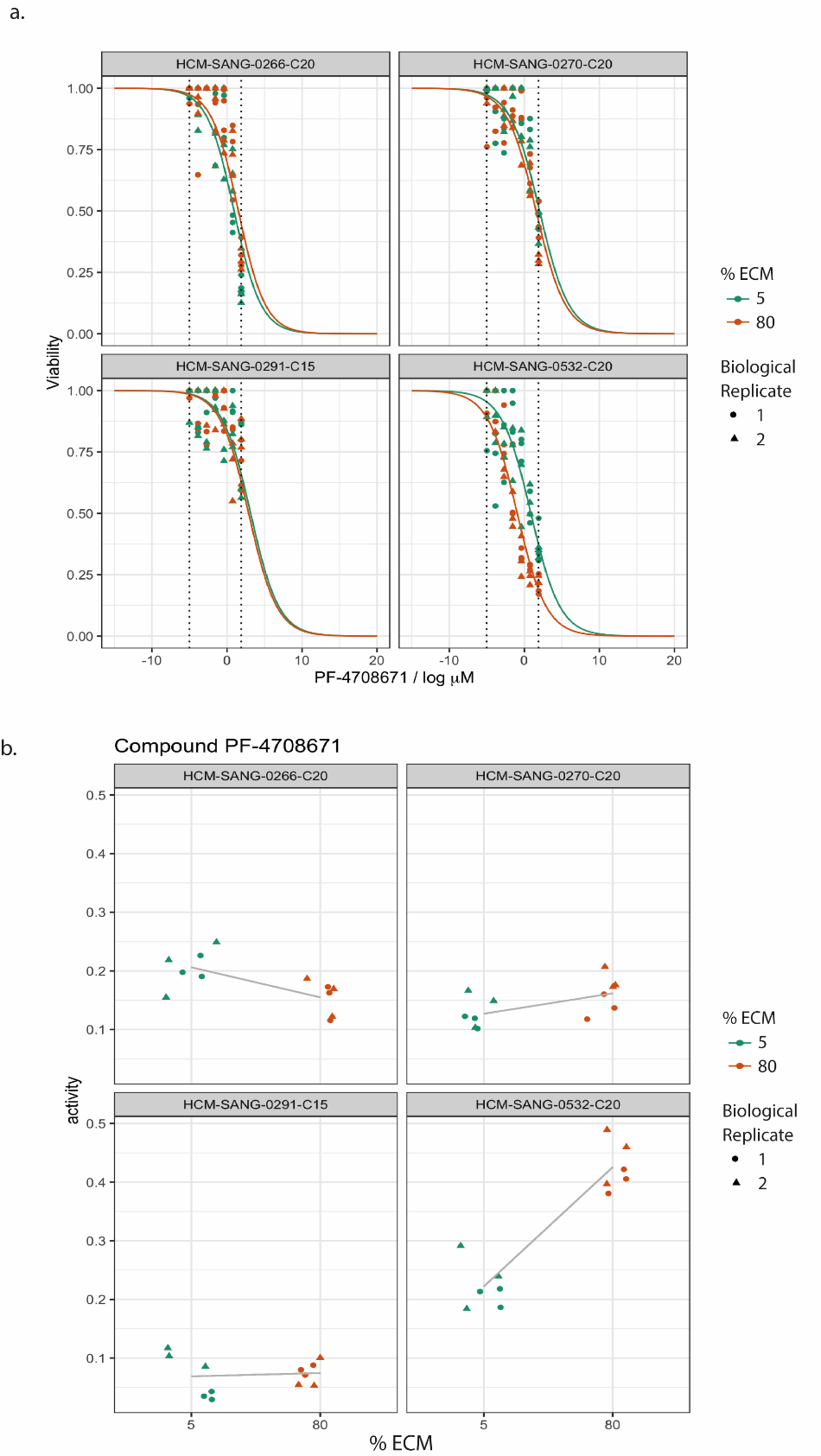

**Figure S5.** *a. Dose response curves showing response of 4 organoid models to PF4708671 in both low 5% ECM and standard 80% ECM conditions, dotted lines indicate the minimum and maximum drug concentration assayed. b. activity plots for the 4 organoid models comparing response to the compound PF4708671 in low 5% ECM and standard 80% ECM conditions.*

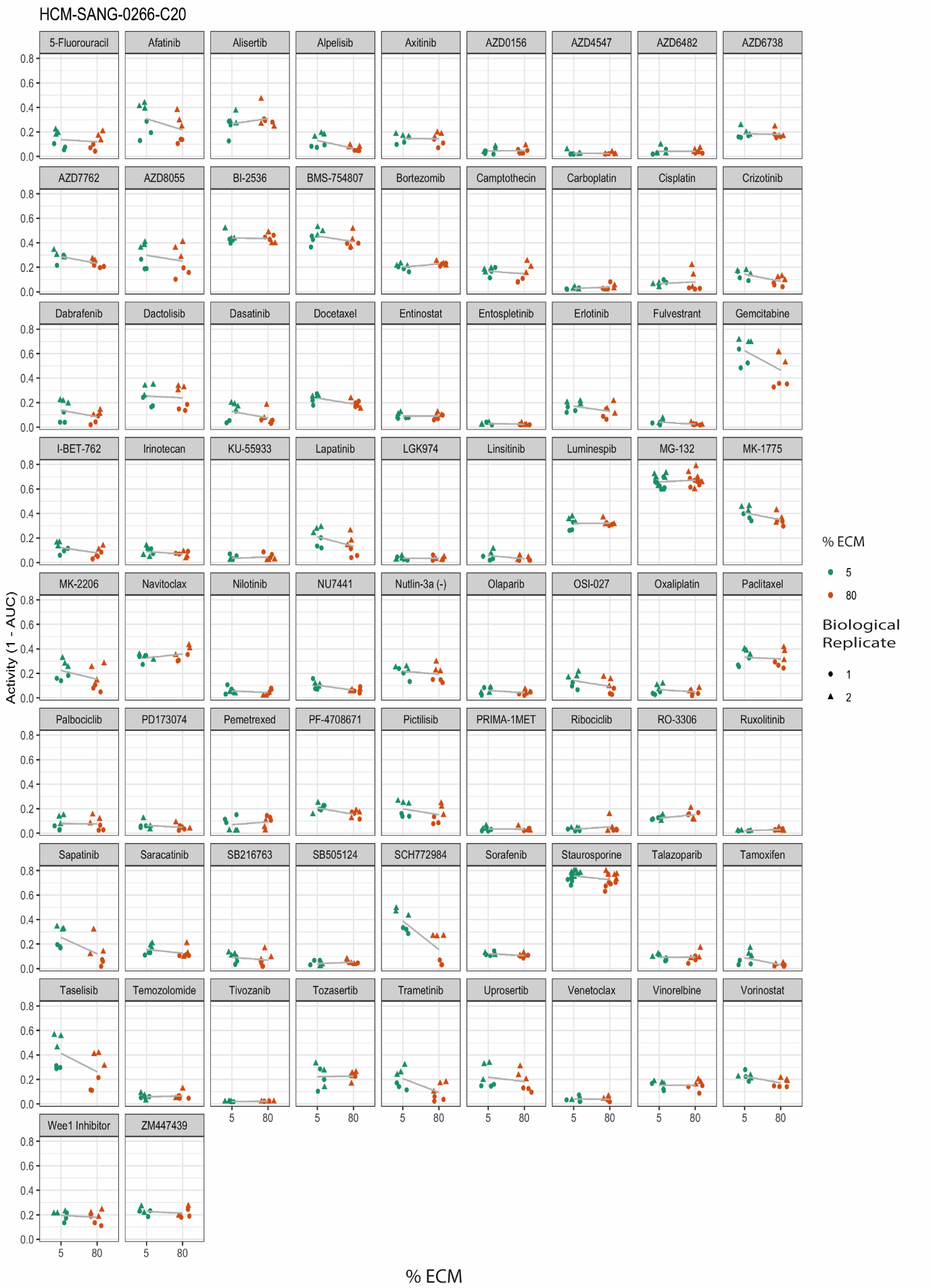
**Figure S6.** *Activity plots for 72 anti-cancer compounds and 2 control compounds (staurosporine and MG132) in HCM-SANG-0266-C20.*

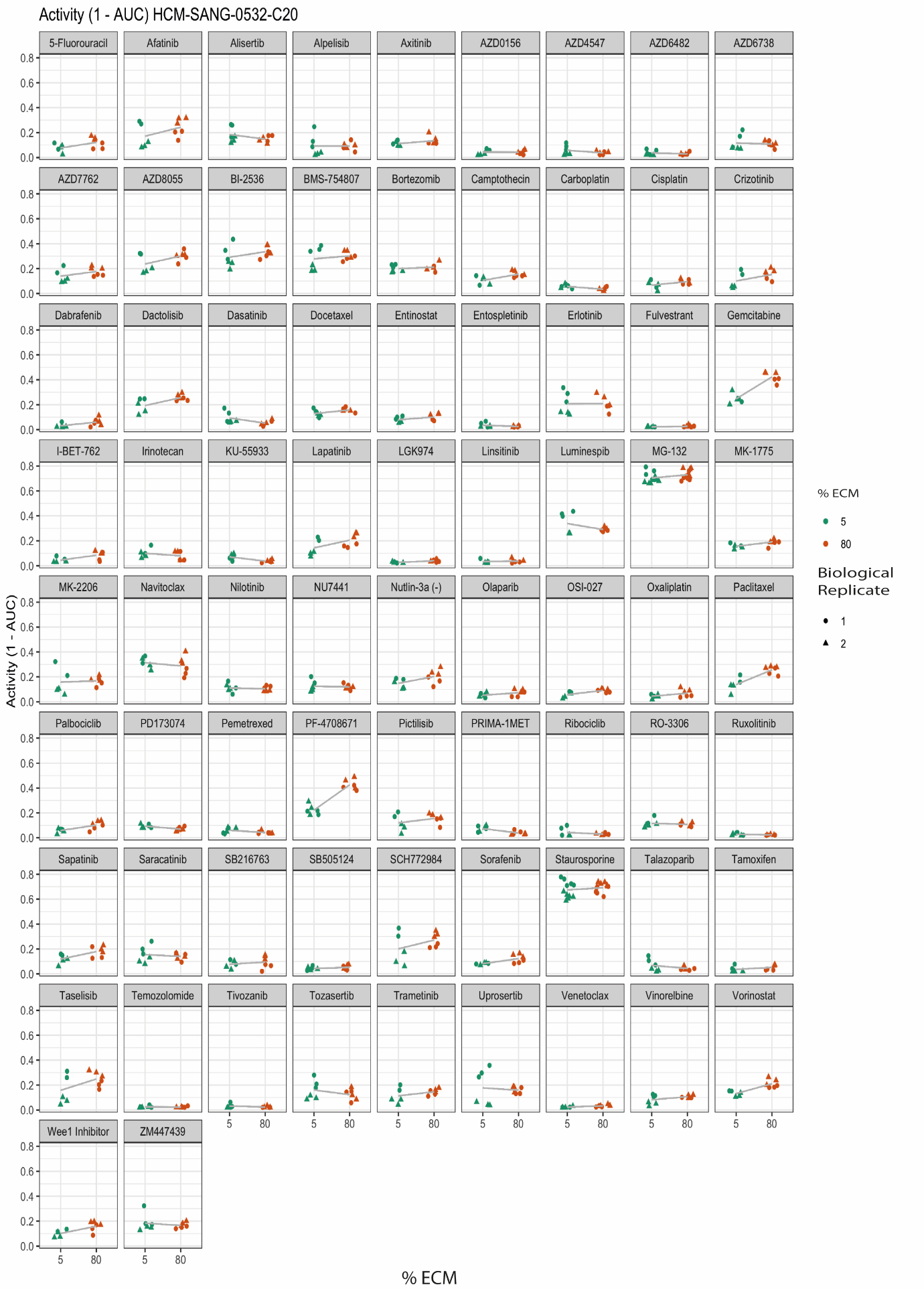

**Figure S7.** *Activity plots for 72 anti-cancer compounds and 2 control compounds in HCM-SANG-0532-C20.*

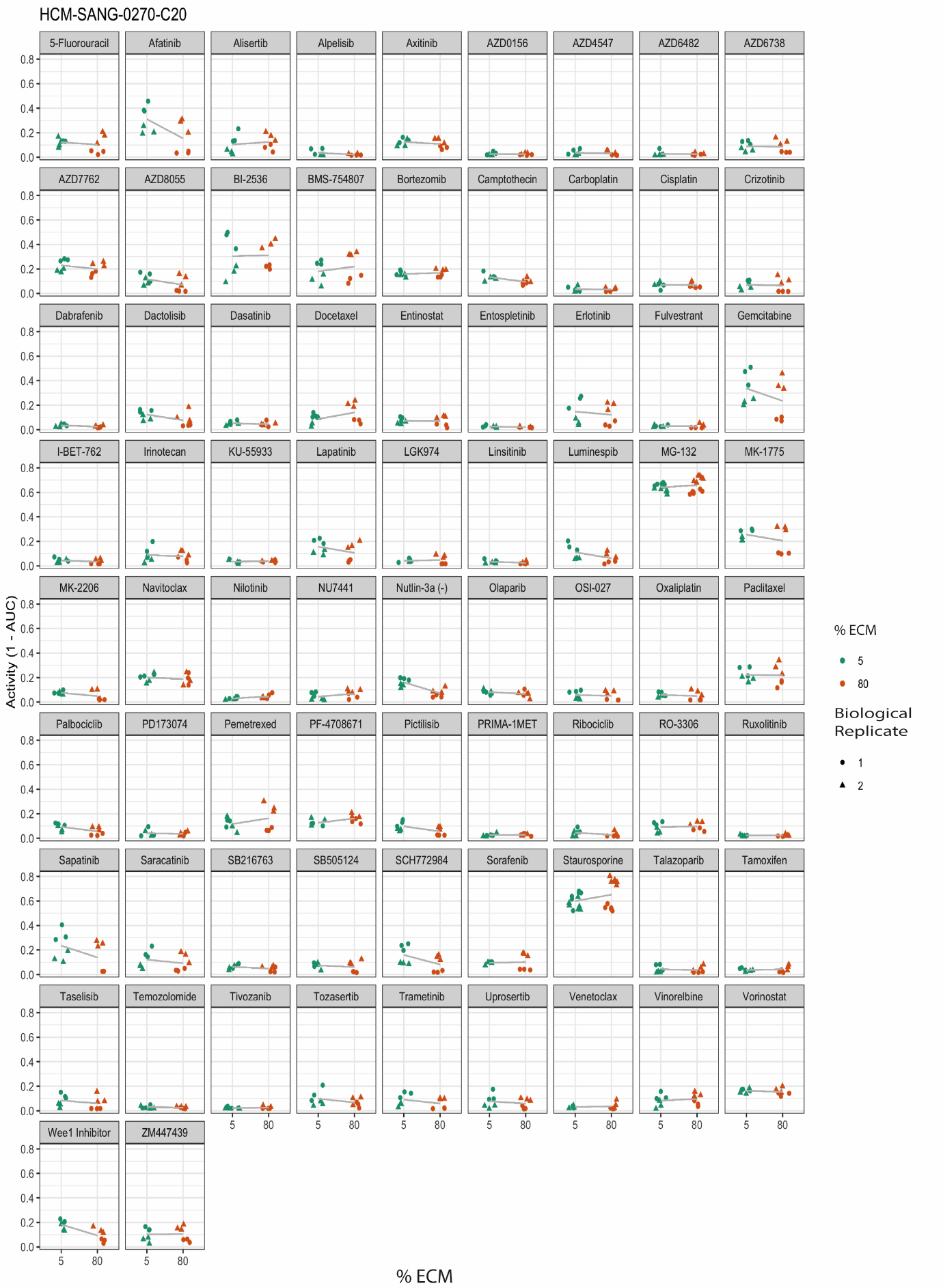

**Figure S8.** *Activity plots for 72 anti-cancer compounds and 2 control compounds in HCM-SANG-0270-C20.*

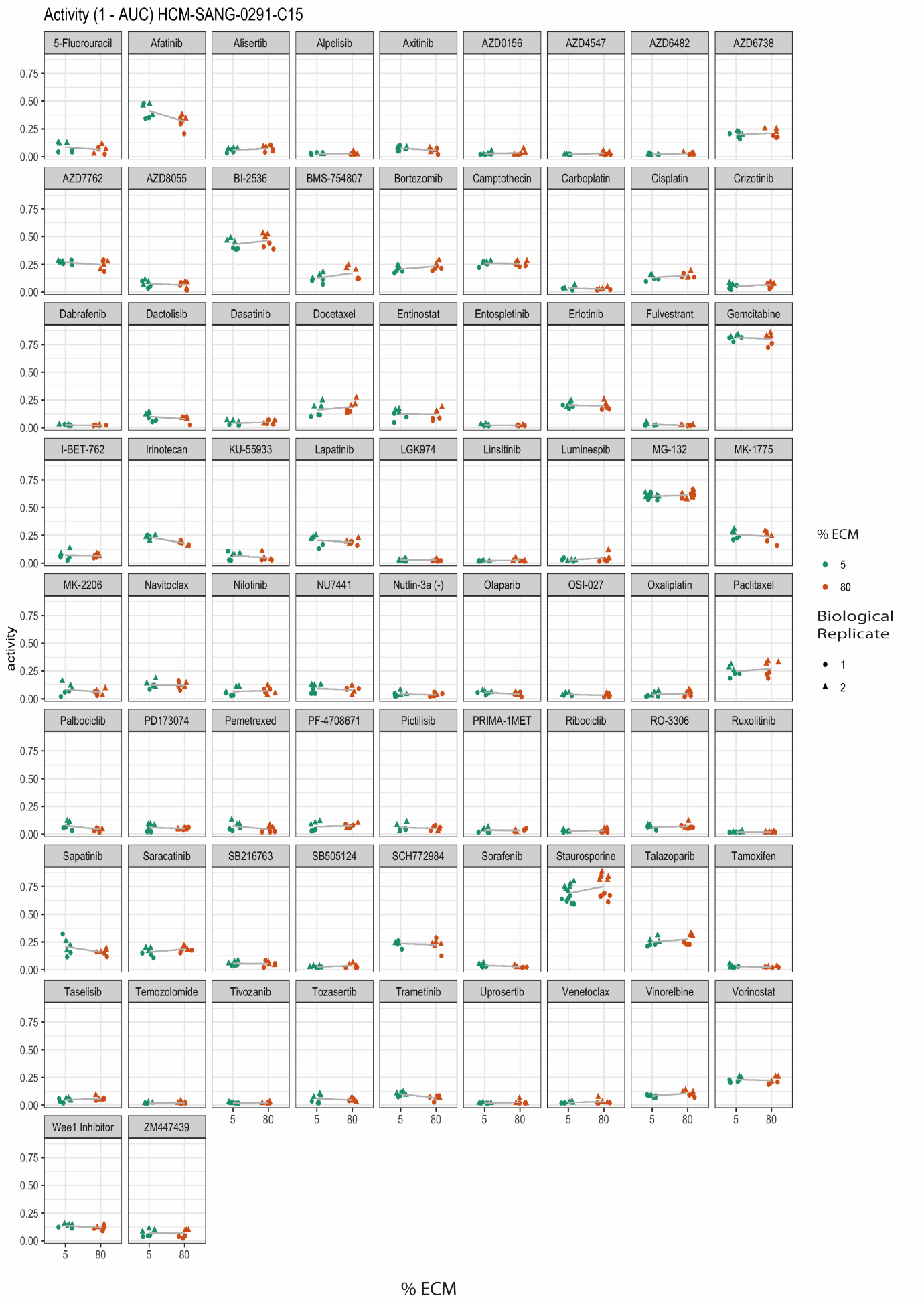
**Figure S9.** *Activity plots for 72 anti-cancer compounds and 2 control compounds in HCM-SANG-0291-C15.*

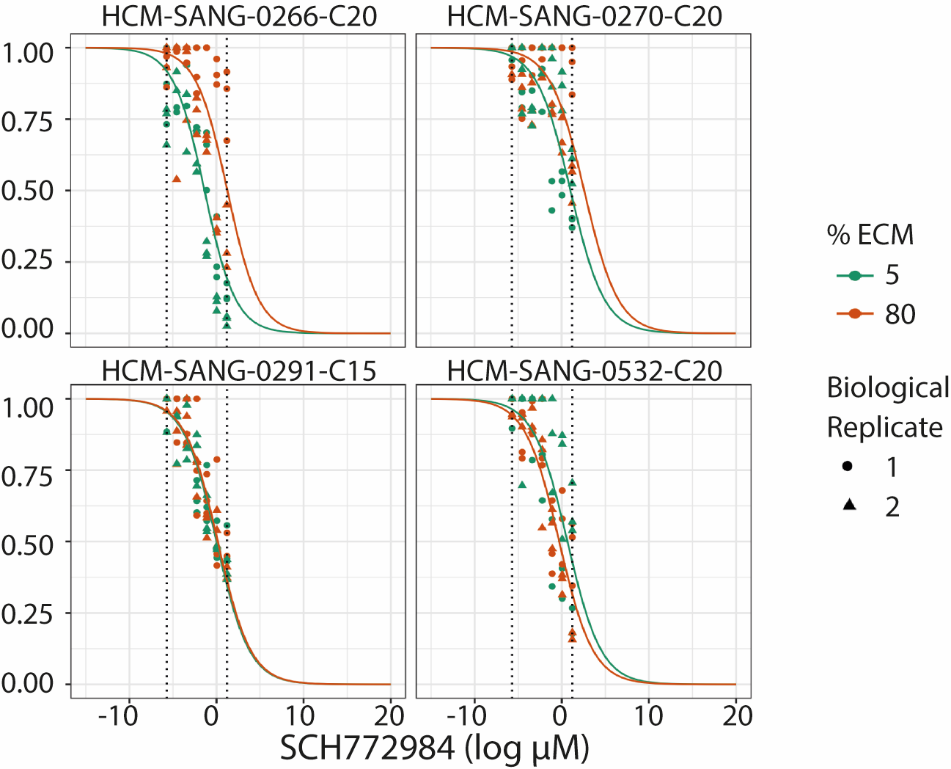

**Figure S10.** *Representative dose response curves for the four organoid models when treated with SCH772984.*

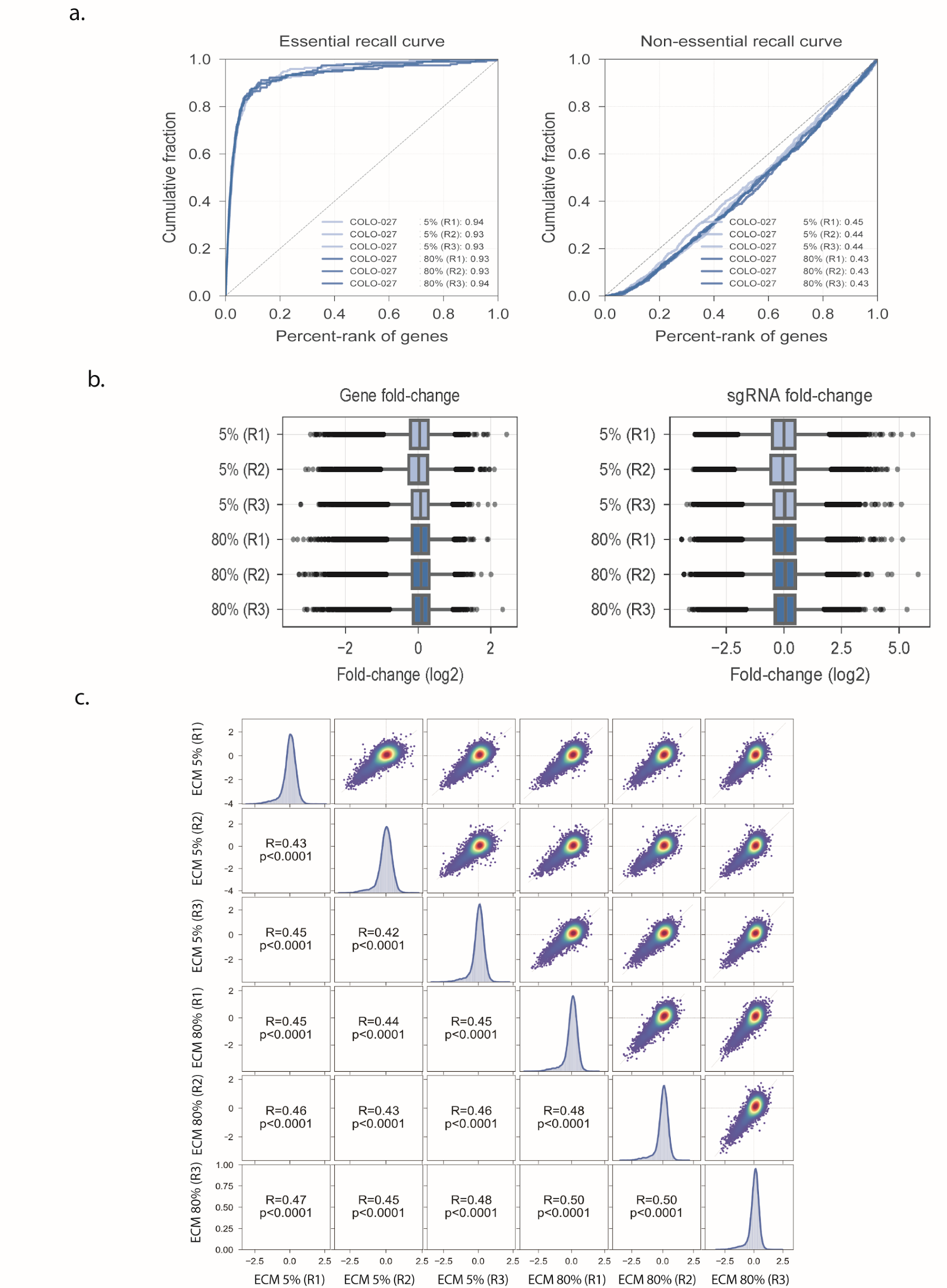

**Figure S11.** *a. Cumulative distribution curves of the HCM-SANG-0273-C18 CRISPR-Cas9 screen performed in 5% ECM and standard 80% ECM conditions for essential genes (left) non-essential genes (right). Three replicates are shown. b. Log_2_ fold changes from library representation at the gene level (left) and gRNA (right) c. Correlation plot of all replicates from arm 1 and arm 2 of the HCM-SANG-0273-C18 CRISPR screen.*
